# Photoacoustic Imaging as a Novel Non-Invasive Biomarker to Assess Intestinal Tissue Oxygenation and Motility in Neonatal Rats

**DOI:** 10.1101/2023.06.27.545971

**Authors:** Victoria G. Weis, Nildris Cruz-Diaz, Jessica L. Rauh, Maryssa A. Ellison, Liliya M. Yamaleyeva, Cherrie D. Welch, Kristen A. Zeller, Jared A. Weis

**Affiliations:** Wake Forest Institute for Regenerative Medicine, Winston-Salem, North Carolina; Department of Surgery-Hypertension, Wake Forest University School of Medicine, Winston-Salem, North Carolina; Cardiovascular Sciences Center, Wake Forest University School of Medicine, Winston-Salem, North Carolina; Department of General Surgery, Section of Pediatric Surgery, Atrium Health Wake Forest Baptist, Winston Salem, North Carolina; Division of Neonatology, Department of Pediatrics, Atrium Health Wake Forest Baptist, Winston-Salem, North Carolina; Department of Biomedical Engineering, Wake Forest University School of Medicine, Winston-Salem, North Carolina; Comprehensive Cancer Center, Atrium Health Wake Forest Baptist, Winston-Salem, North Carolina; School of Biomedical Engineering and Sciences, Virginia Tech-Wake Forest University, Blacksburg, Virginia

**Author notes:** Co-corresponding Authors: Victoria G. Weis, PhD, 391 Technology Way NE, Winston-Salem, NC 27101, Jared A. Weis, PhD, 575 N. Patterson Ave, Ste 530, Winston-Salem, NC 27101. These authors contributed equally (Victoria G. Weis, Jared A. Weis).

**Keywords:** premature infant, intestine, imaging, tissue oxygenation, intestinal motility, necrotizing enterocolitis

## Abstract

**Background:** Within the premature infant intestine, oxygenation and motility play key physiological roles in healthy development and disease such as necrotizing enterocolitis. To date, there are limited techniques to reliably assess these physiological functions that are also clinically feasible for critically ill infants. To address this clinical need, we hypothesized that photoacoustic imaging (PAI) can provide non-invasive assessment of intestinal tissue oxygenation and motility to characterize intestinal physiology and health.

**Methods:** Ultrasound and photoacoustic images were acquired in 2-day and 4-day old neonatal rats. For PAI assessment of intestinal tissue oxygenation, an inspired gas challenge was performed using hypoxic, normoxic, and hyperoxic inspired oxygen (FiO2). For intestinal motility, oral administration of ICG contrast agent was used to compare control animals to an experimental model of loperamide-induced intestinal motility inhibition.

**Results:** PAI demonstrated progressive increases in oxygen saturation (sO2) as FiO2 increased, while the pattern of oxygen localization remained relatively consistent in both 2-day and 4-day old neonatal rats. Analysis of intraluminal ICG contrast enhanced PAI images yielded a map of the motility index in control and loperamide treated rats. From PAI analysis, loperamide significantly inhibited intestinal motility, with a 32.6% decrease in intestinal motility index scores in 4-day old rats.

**Conclusion:** These data establish the feasibility and application of PAI to non-invasively and quantitatively measure intestinal tissue oxygenation and motility. This proof-of-concept study is an important first step in developing and optimizing photoacoustic imaging to provide valuable insight into intestinal health and disease to improve the care of premature infants.

**Highlights:**

- Intestinal tissue oxygenation and intestinal motility are important biomarkers of intestinal physiology in health and disease of premature infants.
- This proof-of-concept preclinical rat study is the first to report application of photoacoustic imaging for the neonatal intestine.
- Photoacoustic imaging is demonstrated as a promising non-invasive diagnostic imaging method for quantifying intestinal tissue oxygenation and intestinal motility in premature infants.

**Graphical abstract:** 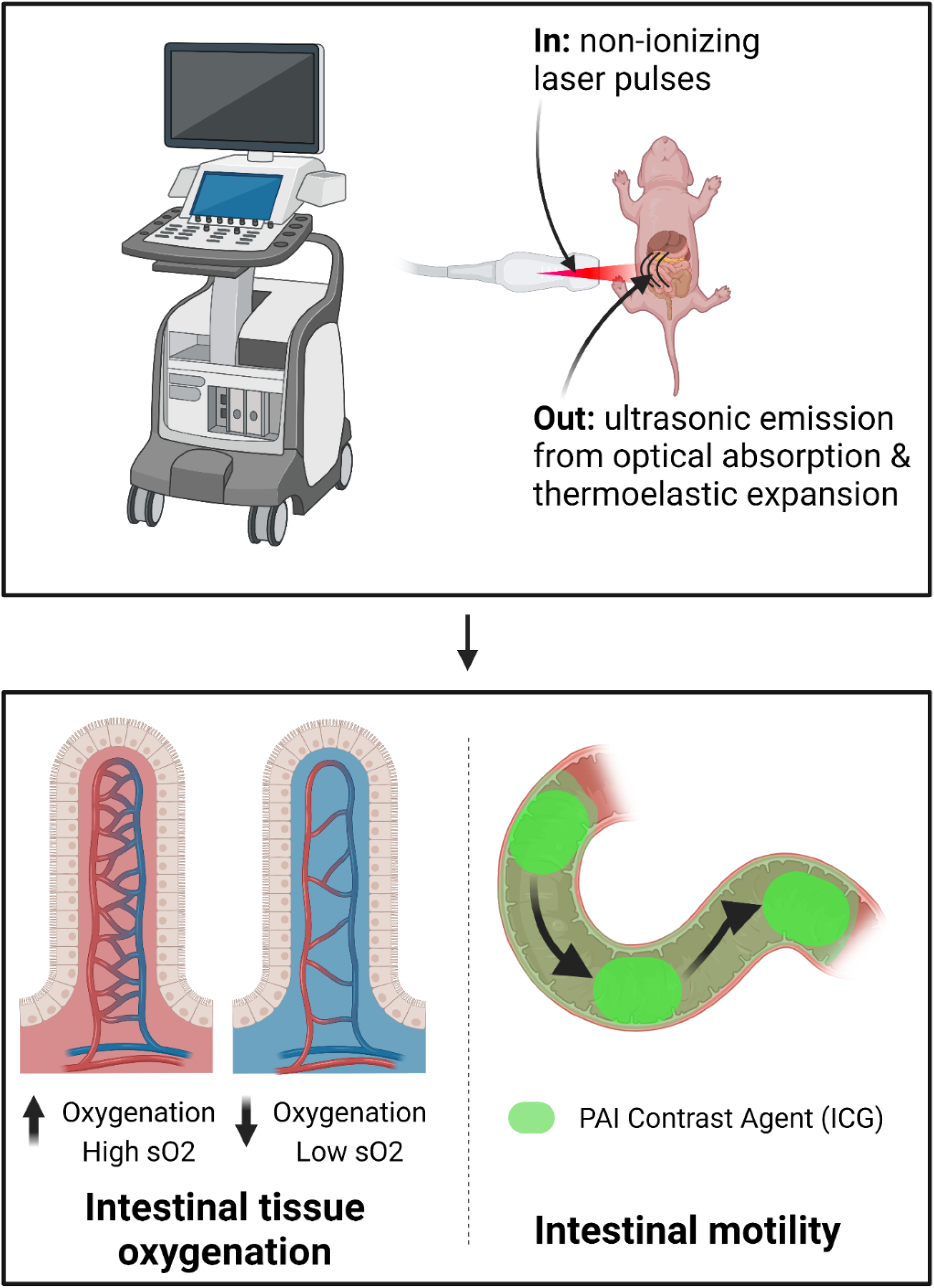

## Introduction

Assessment of intestinal tissue oxygenation and functional motility may play important roles in the overall evaluation of intestinal physiological health and disease in premature infants. Tissue oxygenation is essential for cellular respiration, energy production, and maintenance of tissue integrity. Specifically in development, the preterm infant intestine is particularly vulnerable to hypoxia [1-3]. In hypoxic disease states, the intestinal epithelium experiences oxidative stress, inflammation, and impaired nutrient absorption [4, 5]. Additionally, compromised intestinal tissue oxygenation is strongly linked with the onset and progression of necrotizing enterocolitis (NEC), a life-threatening intestinal disease in premature infants [3, 6, 7]. Thus, assessment of intestinal tissue oxygenation can yield important early diagnostic insights into preterm infant intestinal health and disease, allowing for intervention and management strategies to improve health outcomes in this vulnerable population.

Functional intestinal motility is essential for proper digestion, nutrient absorption, and elimination of waste. Coordinated intestinal peristalsis is essential for movement of nutrients to specialized parts of the gastrointestinal tract. Peristalsis is a multi-factorial process including smooth muscle electrical activity that is influenced by a range of hormonal and neural influences. In intestinal disease states, dysmotility can result in abdominal distension, vomiting or diarrhea, all of which can compromise the absorption of nutrients and elimination of waste products [8, 9]. Additionally, dysmotility can alter the gut microbiome and further perpetuate dysbiosis [10]. Assessment of abnormality and dysfunction in intestinal motility is critical for understanding the underlying pathophysiology to adequately identify and treat infants with intestinal dysmotility [11].

To date, accurate and conclusive early diagnosis of premature infant intestinal pathologies such as necrotizing enterocolitis remains elusive. Current diagnosis relies on coarse assessment methods based on combinations of clinical signs and symptoms and radiographic features that are often nonspecific. Advanced non-invasive imaging modalities that may provide increased diagnostic precision for the intestinal pathologies (e.g., MRI, CT) present significant challenges in infants, especially in the medically complex NICU patient population. Specifically, radiation exposure from CTs is of concern, and in most centers, infants must be transported out of the NICU to these scanners located elsewhere in the hospital, placing the infant at risk of not having the appropriately trained personnel and equipment available in the event of a clinical emergency. Additionally, preterm infants are unable to undergo the prolonged MRI scan as they will become profoundly hypothermic outside of their incubator. Other non-invasive imaging methods such as ultrasound are being developed but have yet to be validated and fully effective for assessment of premature infant intestinal pathologies. Thus, serial plain-film abdominal X-ray (XR) examination remains the clinical standard for diagnosis. One of the most serious infant gastrointestinal diseases is necrotizing enterocolitis (NEC), where radiographic diagnosis is made upon detection of intestinal wall pneumatosis, with or without pneumoperitoneum [12, 13]. By the time this finding is seen on serial XRs, it may have progressed to an advanced stage that necessitates urgent surgical management with associated high morbidity and mortality [12]. Bowel ultrasound imaging is an emerging technique to improve diagnosis with high diagnostic specificity. However, ultrasound imaging for NEC detection, as well as delineating other intestinal pathologies, has not yet gained wide clinical adoption, primarily due to challenges in interpretation and diagnostic uncertainty with demonstrated poor sensitivity and an estimated false negative rate of up to 40% [14]. New accurate and clinically translatable diagnostic methods for quantitative assessment of intestinal tissue oxygenation and functional motility are urgently needed to allow for effective disease diagnosis and management in developing infants.

Photoacoustic imaging (PAI) is an emerging non-invasive imaging modality with high spatial and temporal resolution. PAI takes advantage of the ‘optoacoustic effect’ to generate image contrast through ultrasound detection of optical absorption events [15-17]. Pulses of near-infrared light are absorbed by tissue chromophores which undergo thermoelastic expansion and produce acoustic signal. This acoustic signal production is then detected by ultrasound transducers. This hybrid approach of ‘light in – sound out’ provides more sensitive/specific detection than ultrasound imaging and significantly increased penetration depth over optical imaging [15]. As hemoglobin is a strong optical absorber, the direct PAI detection of optical absorption in hemoglobin can be used as basis for quantitative image contrast. Hemoglobin exhibits distinctly different absorption spectra based on its oxygenation state. By incorporating spectrally selective excitation and imaging, oxy- and deoxyhemoglobin can be separately visualized and quantified to allow for assessment of spatial tissue oxygen saturation. Additionally, exogenous small molecule dyes with near-infrared optical absorption (e.g., FDA-approved indocyanine green dye) can be used as exogenous contrast agents to enhance PAI signal intensity.

PAI has been investigated for a range of clinical disease-site applications including breast, skin, peripheral vascular diseases, musculoskeletal, gastrointestinal, and adipose tissue [15]. However clinical translation of PAI technologies has been limited due to fundamental physical limitations in penetration depth of less than ∼ 7 - 10 cm [15, 17]. While methods to increase the penetration depth of PAI and accuracy of PAI measurements in deeper tissues are actively being explored to advance this non-invasive imaging technology, the application of PAI to the developing infant presents an especially promising setting as penetration depths in commercially available PAI systems are capable of imaging throughout the body, especially in premature infants at highest risk for disease. PAI also has particular appeal for use in imaging premature infants as there is no ionizing radiation, patient transport, or sedation required. PAI is therefore well positioned to assess complicated intestinal disease diagnostic features, specifically physiological hallmarks of intestinal disease in the premature infant for which current clinical diagnostic tools are inadequate or infeasible, including intestinal oxygenation and dysmotility.

In this study, we hypothesized that PAI would provide a novel non-invasive quantitative assessment of intestinal tissue oxygenation and motility to characterize changes in intestinal physiology in the neonatal rat. This study represents, to our knowledge, the first use of PAI to assess the developing intestine of neonates and infants. We developed imaging acquisition and image analysis methods for the use of photoacoustic imaging to assess physiological indicators of intestinal health based on assessment of intestinal tissue oxygenation and intestinal motility in neonatal rat pups. We performed several validation studies with experimental perturbations known to modulate intestinal tissue oxygenation and intestinal motility, respectively, as a proof-of-concept validation of our imaging and analysis methods.

## Methods

### Animals

The care, maintenance, and treatment of animals in these studies adheres to the protocols approved by the Institutional Animal Care and Use Committee of Wake Forest University. Timed-pregnant Sprague-Dawley rats were obtained from Charles River Laboratories (Wilmington, MA, USA) and monitored for birth beginning 36□ s before expected birth. Date hour of birth was recorded, and pups remained with their dams for the duration of studies. Rat pups were used at either 2-day and 4-day old for all studies. Upon completion of imaging sessions, pups were returned to dams and no infanticide incidences occurred.

### Photoacoustic imaging

Ultrasound and photoacoustic images were acquired using a VEVO 2100 LAZR High-Resolution Ultrasound and Photoacoustic Imaging System (FUJIFILM VisualSonics, Inc., Toronto, ON, Canada). Inhalation anesthesia was induced in all animals via a 2%/98% isoflurane/medical air mixture and maintained via a 1%/99% isoflurane/medical air (or hypoxic/hyperoxic gas as noted in experimental conditions) mixture. Animals were placed supine on a heated imaging stage with continuous physiological monitoring. An LZ-550 transducer with axial resolution of 44μm and broadband frequency of 32 MHz - 55 MHz was used to image in the sagittal plane with centrifuged ultrasound gel to provide acoustic coupling. All imaging protocols were standardized with consistent US/PAI settings (40 MHz Tx Frequency, 100% 2D Power, 100% PA Power, 6 Hz frame rate, 40 dB PA gain, 18 dB 2D gain, 12 mm depth, 14mm width, 20% threshold hemoglobin) and imaging depth across all animals.

For imaging tissue oxygenation, PAI was acquired in the sagittal plane at the abdominal midplane in each animal in Oxy-Hemo mode to obtain maps of tissue oxygen saturation (sO2) based on automated dual wavelength PAI excitation at 750 nm and 850 nm. Calculation of sO2 is automatic within the VEVO LAZR system and is based on the ratio of oxyhemoglobin to total hemoglobin. sO2 maps were recorded for subsequent oxygenation analysis. Volumetric maps of sO2 were similarly acquired using a motorized ultrasound probe positioner with slice thickness of 0.3 mm and 50 total slices covering the entire abdominal area.

For imaging intestinal motility, 7.5 mg/kg indocyanine green (ICG) (Indocyanine Green for Injection, Diagnostic Green, Farmington Hills, MI, USA) diluted in sterile water was administered via oral gavage in rat pups approximately 1.5 hours prior to the imaging session. PAI was acquired in the sagittal plane at the abdominal midplane in each animal in single spectra excitation mode at 810 nm wavelength. 810 nm was selected to coincide with the peak of the ICG absorption spectra when dissolved in water [18]. Cine time series recordings covering a minimum of 30 seconds (at 6 Hz acquisition frame rate) of intestinal peristalsis were recorded for subsequent motility analysis.

### Intestinal tissue oxygenation assessment

To optimize and validate PAI measurement of intestinal tissue oxygenation, we performed a proof-of-concept study using an inspired gas challenge using hypoxic, normoxic, and hyperoxic gas with fractions of inspired oxygen (FiO2) of 5%, 21%, and 100%, respectively. Hypoxic gas was provided from a 5% oxygen/95% nitrogen compressed gas cylinder. Normoxic gas was provided from a medical air compressed gas cylinder. Hyperoxic gas was provided from a 100% oxygen compressed gas cylinder. Compressed gas cylinders (Linde Gas & Equipment Inc, Burr Ridge, IL) were connected to the inhalation anesthesia vaporizer with 1 L/min flow rate. Imaging sessions were started under normoxic conditions. Following completion of image acquisition, the inhaled gas was manually changed via a flow control valve to the hypoxic gas and allowed to equilibrate for 5 minutes before image acquisition. Following completion of image acquisition, the inhaled gas was then manually changed via a flow control valve to the hyperoxic gas and allowed to equilibrate for 5 minutes before image acquisition.

In cohorts of 2-day and 4-day old neonatal rats (n = 5 animals at each time point), animals were imaged using the VEVO 2100 LAZR PAI system at each FiO2 level (normoxic, hypoxic, hyperoxic) and tissue oxygenation was measured using the VEVO Oxy-Hemo imaging mode which uses spectral wavelengths of 750 nm and 850 nm to calculate the ratio of oxygenated hemoglobin concentration to total hemoglobin concentration based on PAI signal intensity. A region-of-interest (ROI) was manually defined over the intestinal region within the abdomen and an average value of tissue oxygen saturation (sO2) within this intestinal ROI was extracted as a measure of intestinal tissue oxygenation. sO2_Average_ values were recorded in this analysis, which represents the average oxygen saturation within the ROI after excluding values of zero. This post-processing of the Oxy-Hemo images was performed using the OxyZated tools in the Vevo LAB postprocessing software (FUJIFILM VisualSonics, Inc., Toronto, ON, Canada).

### Intestinal motility assessment

To optimize and validate PAI measurement of intestinal motility, we performed a proof-of-concept study comparing control animals to an experimental model of drug-induced inhibition of intestinal motility via acute loperamide administration. In a cohort of 4-day old neonatal rats (n = 3 animals at each time point), animals were administered loperamide hydrochloride (Medline, Northfield, IL, USA) at 10 mg/kg diluted in sterile water via oral gavage to inhibit intestinal peristalsis or vehicle control (sterile water) two hours prior to the imaging session. At 1.5 hours prior to the imaging session, ICG contrast agent was administered to both cohorts as an intraluminal contrast agent. To assess intestinal peristalsis, ∼30 seconds cine recordings of US and PAI signal intensity were recorded using the VEVO 2100 LAZR PAI system with PAI spectral excitation at the ICG peak wavelength (810 nm).

As shown in Figure 1, post-processing of both US and ICG-weighted PAI images was used to assess intestinal motility. Frame-by-frame motion deformation estimates of intestinal peristalsis throughout the cine time course were extracted using a deformable symmetric normalization method (SyN) diffeomorphic nonrigid image registration available within the ANTs repository [19]. Each image in the cine series was registered to the initial image using an initial rigid/affine registration followed by the deformable ANTs SyN nonrigid registration method. Parameters for ANTs SyN were as follows: gradient step size: 0.1, spline distance: 26, histogram matching: off, cross correlation radius: 4. Following deformation estimation, we calculated a map of the “intestinal motility index” from US and PAI images based on an analysis method previously established for assessment of small bowel motility from clinical dynamic MR images [20]. To calculate this motility index, pixel-wise intestinal deformation estimates were first used to construct a Jacobian matrix of the deformation field containing the spatial gradients of the deformation vector at each pixel location. The determinant of the deformation Jacobian matrix was then calculated as an index to quantify the volumetric change of each pixel at each time frame within the cine series as an estimate of expansion/contraction due to intestinal peristalsis. This Jacobian determinant of the deformation field is a commonly used metric to assess deformation in biological tissue as it is insensitive to global rigid motion (e.g., respiratory motion) with low values indicating low/no expansion/contraction and high values indicating high expansion/contraction. This analysis yields a quantitative index of intestinal motility for each frame within the cine time series. The standard deviation across the time dimension was then used to reduce dimensionality to generate an overall summary image of the intestinal motility index quantifying variance of intestinal motility across the entire cine recording. An ROI was then manually drawn over the intestinal area within the abdomen based on the anatomical US image. The mean value of the motility index (standard deviation across time of the Jacobian determinant of the deformation field) within the ROI was used as a summary score of the intestinal motility index. This analysis was performed on deformation fields estimated from both anatomical US and ICG-weighted PAI cine images for each animal in each cohort.

**Figure 1.**
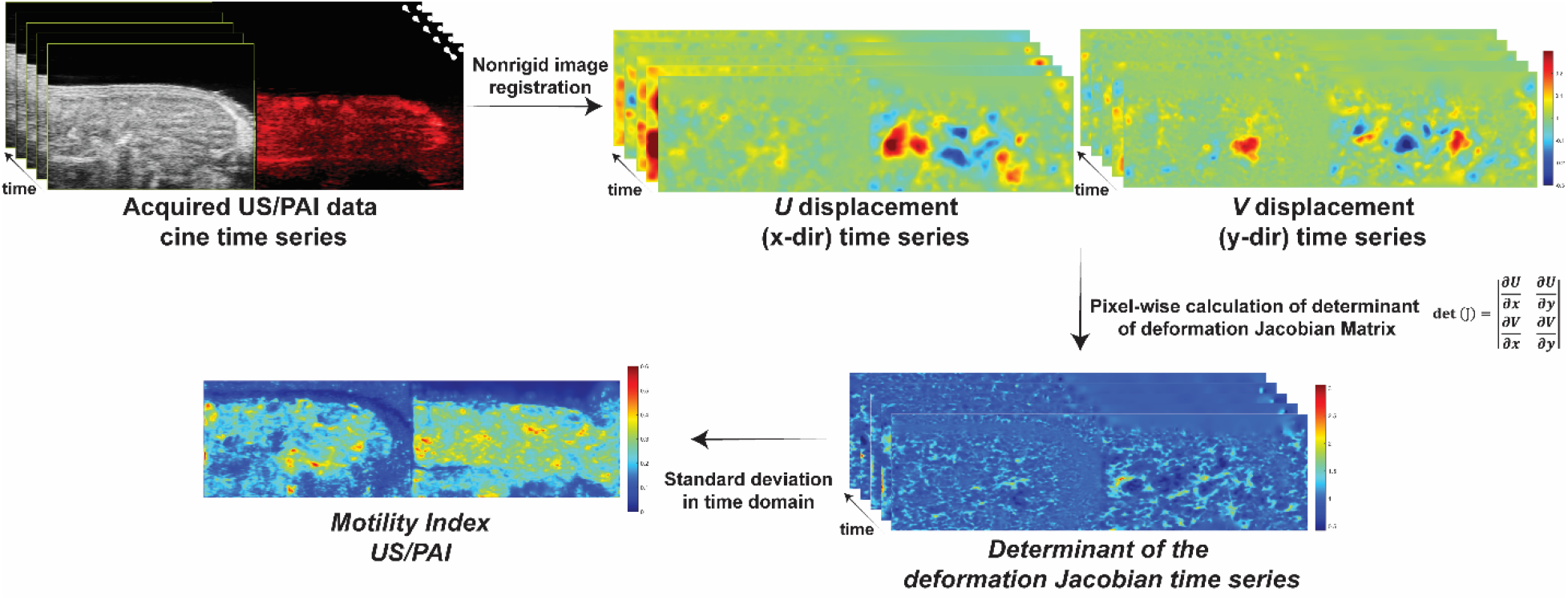
Schematic of the intestinal motility index calculation. Acquired cine time series images from US and PAI imaging of the neonatal rat intestine are registered to the initial image using a nonrigid image registration algorithm to calculate displacements in the x- and y-directions for each image in the time series. Spatial gradients of the deformation fields are then used to create a deformation Jacobian matrix at each pixel location and the determinant of this matrix is used to quantify volumetric expansion/contraction for each pixel location for each image in the time series. The motility index is then calculated based on the standard deviation across the time series.

### Intestinal transit assay

To validate PAI measures of intestinal motility, we performed a small intestinal transit assay in control and loperamide treated animals. Animals were administered 100 uL of a methylene blue 2% aqueous solution, diluted in sterile water, via oral gavage. Dye was allowed to transit for 1 hour and animals were then sacrificed via cervical dislocation. The GI tract from stomach to distal colon was excised and photographed to assess the visibility of methylene blue along the small intestine. Images were analyzed for small intestinal transit by measuring the distance from the stomach-duodenal junction to the most distal location of visibly detected methylene blue. Distances were calibrated using a ruler that was placed within the image field-of-view. Distances were recorded as a small intestinal transit velocity (transit velocity = maximal dye transit distance / 1 hour transit time).

### Statistical analysis

For intestinal tissue oxygenation studies, mean tissue oxygenation within the intestinal ROI at 5%, 21%, and 100% FiO2 was compared groupwise between FiO2 levels in both 2-day and 4-day old animals using a repeated measures one-way ANOVA followed by Holm-Sidak’s multiple comparisons testing between each FiO2 level. For intestinal motility studies, mean motility index within the intestinal ROI in control and loperamide treated animals was compared groupwise using an unpaired t-test. All analyses were completed using GraphPad Prism software with *p*- value < 0.05 considered as statistically significant.

## Results

### Photoacoustic imaging of intestinal tissue oxygenation reveals changes in response to perturbations in inspired gas oxygen fraction

Representative images from PAI assessment of tissue oxygenation for each FiO2 level within the inspired gas challenge are shown in Figure 2 and Figure 3 for 2-day and 4-day old rat pups, respectively. PAI images qualitatively demonstrated progressive increases in sO2 as the fraction of oxygen within the breathing gas increased. While the overall sO2 levels increased, the localization pattern of the oxygen saturation remained relatively consistent throughout the inspired gas challenge. Volumetric renderings of sO2 shown in Figure 4, further demonstrated progressively increasing oxygenation throughout all regions of the small intestine in response to perturbations of increasing fraction of oxygen in inspired gas. Figures 2 and 3 demonstrate quantitative changes in mean intestinal sO2 for each FiO2 level in the inspired gas challenge. In 2-day old rat pups, sO2 increased from 51.13% (+/- 3.84), to 55.90% (+/- 3.77), and 62.58% (+/- 2.19) in 5%, 21%, and 100% FiO2, respectively. These sO2 increases indicated a 9.3% increase in intestinal tissue sO2 from hypoxic breathing gas to normoxic breathing gas and a 11.9% increase in intestinal tissue sO2 from normoxic breathing gas to hyperoxic breathing gas. In 4-day old rat pups, sO2 increased from 49.62% (+/- 3.42) to 56.13% (+/- 3.52), and 69.02% (+/- 3.47) in 5%, 21%, and 100% FiO2, respectively. This resulted in a 13.1% increase in intestinal tissue sO2 from hypoxic breathing gas to normoxic breathing gas and a 23.0% increase in intestinal tissue sO2 from normoxic breathing gas to hyperoxic breathing gas. Differences in intestinal tissue sO2 at each inspired gas FiO2 level (hypoxic vs. normoxic, normoxic vs. hyperoxic, and hypoxic vs. hyperoxic) were statistically significant within both 2- day and 4-day old rat pups. Tissue oxygenation levels (sO2) and tissue reactivity (changes in sO2 between FiO2 levels) within the intestine can vary depending on age and development stage. Thus, sO2 levels and reactivity changes between FiO2 levels were also compared between 2-day and 4-day old animals. With 100% FiO2, sO2 was significantly increased 10.3% in 4-day old animals compared to 2-day old animals (4-day: 69.02% +/- 3.47 vs 2-day 62.58% +/- 2.19, p<0.01). No statistically significant differences in intestinal tissue sO2 were found between 2-day and 4-day old age groups at hypoxic and normoxic breathing gas settings. Similarly, no difference in tissue reactivity to changes in FiO2 (normoxic to hypoxic or normoxic to hyperoxic) was observed in 2-day compared to 4-day old age groups.

**Figure 2.**
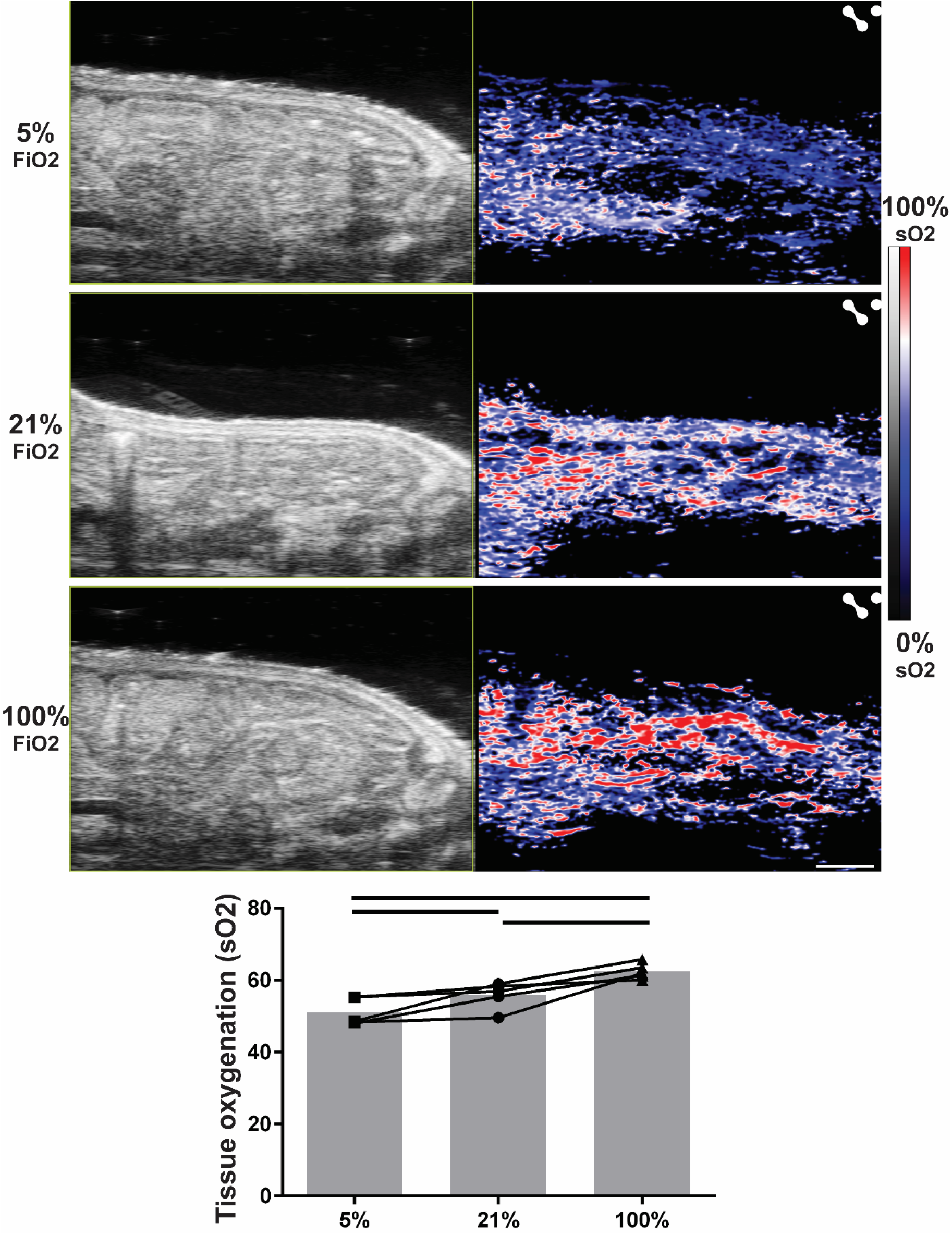
Intestinal oxygen saturation in 2-day old pups at 5%, 21%, and 100% FiO2. Progressive increases in intestinal tissue oxygenation (blue=low oxygen saturation, red=high oxygen saturation) are seen when increasing the fraction of inspired oxygen. Graphs depict quantification of intestinal tissue oxygenation in 2-day old rat pups at 5%, 21%, and 100% FiO2 with bars indicating statistical significance at the *p* < 0.05 level.

**Figure 3.**
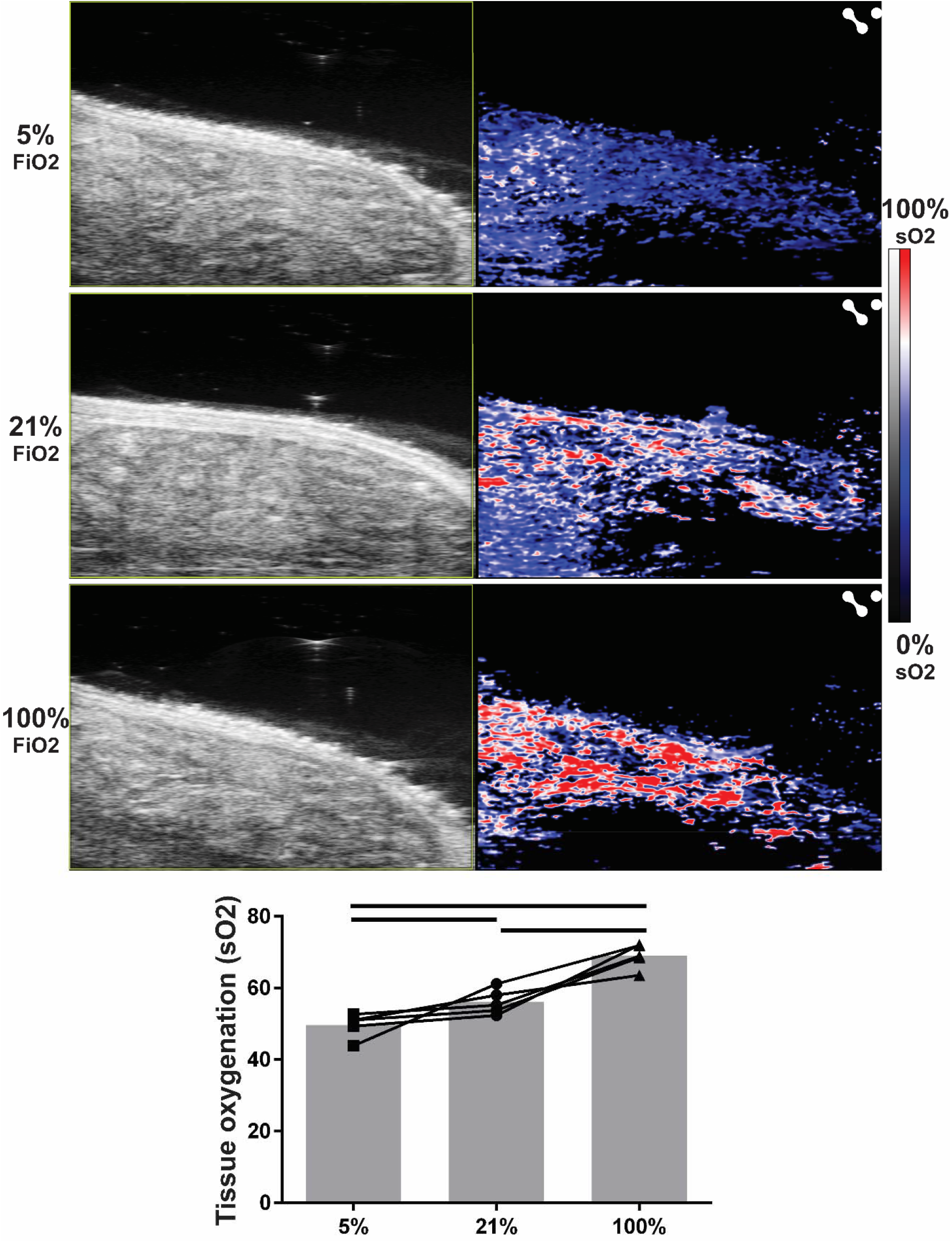
Intestinal oxygen saturation in 4-day old pups at 5%, 21%, and 100% FiO2. Progressive increases in intestinal tissue oxygenation (blue=low oxygen saturation, red=high oxygen saturation) are seen when increasing the fraction of inspired oxygen. Graphs depict quantification of intestinal tissue oxygenation in 4-day old rat pups at 5%, 21%, and 100% FiO2 with bars indicating statistical significance at the *p* < 0.05 level.

**Figure 4.**
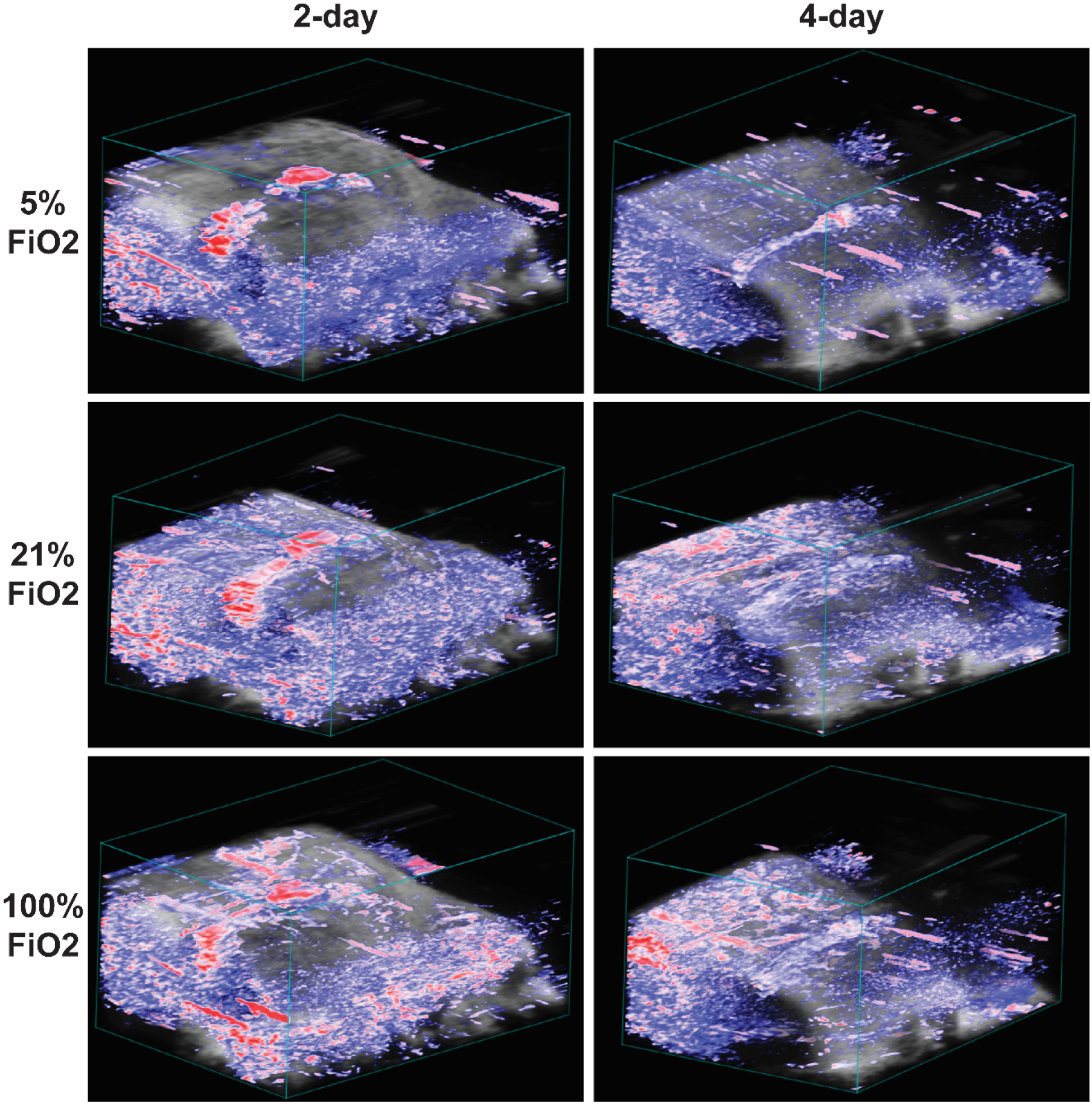
Volumetric renderings of intestinal oxygen saturation in 2-day and 4-day old pups at 5%, 21%, and 100% FiO2. Volumetric renderings will allow for quantitative analysis throughout all intestinal regions.

### Photoacoustic imaging reveals intestinal motility changes in response to acute drug-induced inhibition of intestinal peristalsis

Representative images of US and intraluminal ICG contrast enhanced PAI imaging along with post-processed motility index images from both US and PAI are shown in Figure 5 for 4-day old rat pups in both control and loperamide treated groups. The US and PAI images within these figures represent a single time frame extracted from a cine time series covering ∼30 seconds (at 6 Hz acquisition frame rate) of intestinal peristalsis. Movies for each of the representative images are included within the supplementary materials as Supplemental Movie 1 and 2. The intestinal motility index was calculated for both US and PAI imaging. From US image analysis, the intestinal motility index was decreased by 21.5% in loperamide-treated animals compared to control animals at 4-days old (Control: 0.23 +/- 0.016 a.u. to Loperamide: 0.18 +/- 0.0041 a.u., p<0.01). The use of the intraluminal ICG contrast agent was well-tolerated and yielded an increase in resolution sensitivity with significant intestinal PAI signal intensity at the ICG- weighed 810 nm excitation wavelength. A representative image showing intestinal PAI signal intensity at the ICG-weighted 810 nm excitation wavelength in a 4-day old rat pup without ICG contrast agent is shown in Supplemental Figure 1. As expected, loperamide was found to significantly inhibit intestinal motility, with intestinal motility index scores decreasing 32.6% (Control: 0.26 +/- 0.0092 a.u. to Loperamide: 0.17 +/- 0.0024 a.u., p<0.001) when comparing 4- day old control animals to loperamide-treated animals. Due to differences in signal origins between US and ICG-PAI, the intestinal motility indexes can not be directly compared between the imaging modalities.

**Figure 5.**
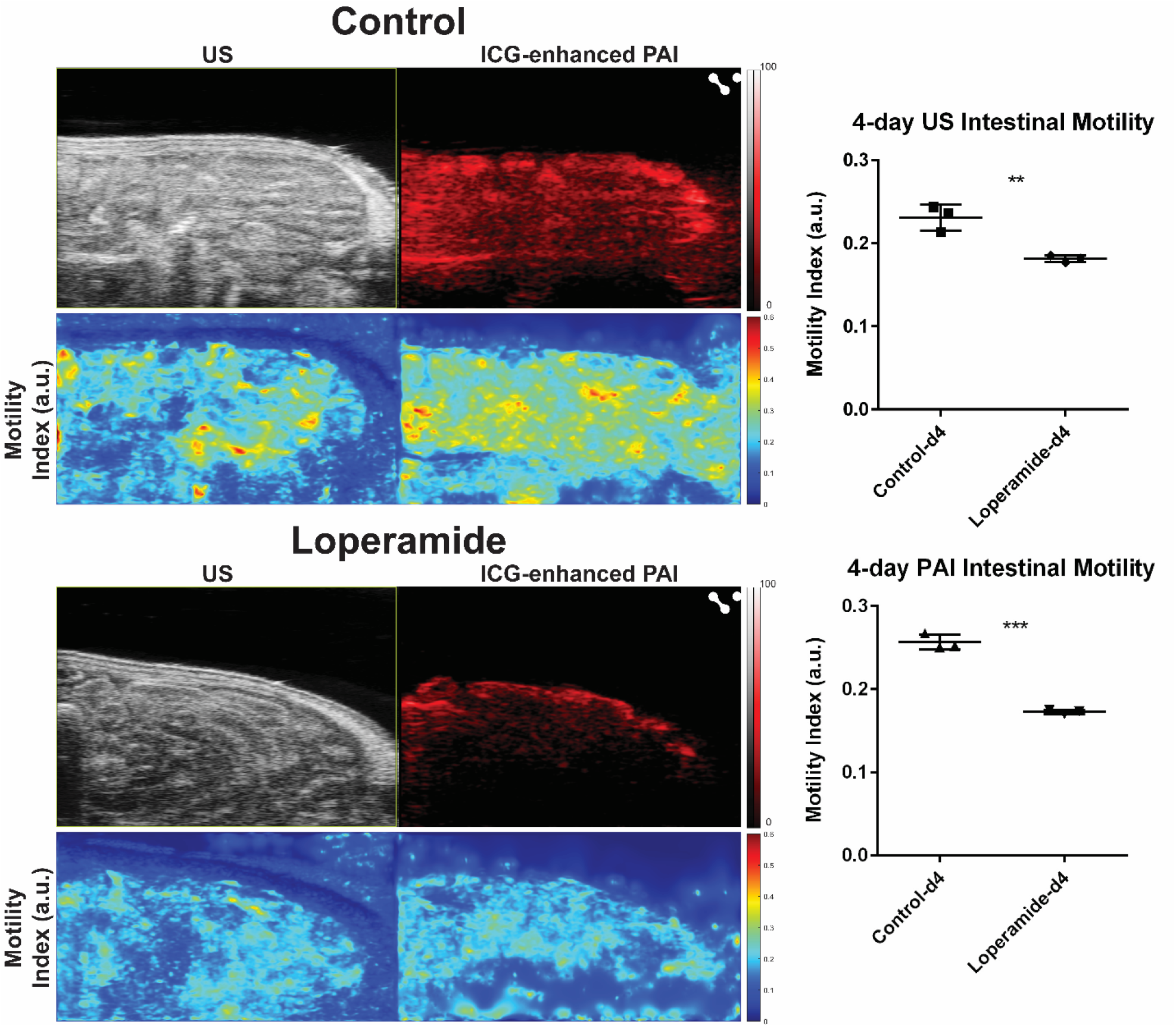
Intestinal motility in 4-day old control and loperamide-treated animals. Left panels are representative images from 4-day old animals. A significant global reduction in the intestinal motility index is seen in loperamide-treated animals as compared to controls. Graphs depict quantification of intestinal motility index in 4-day old rat pups in the control and loperamide-treated groups using both US and PAI images. (**denotes p < 0.01, *** denotes p < 0.001).

### Intestinal transit assay confirms reduction of intestinal motility in response to acute drug-induced inhibition of intestinal peristalsis

Representative images and analysis from the intestinal transit assay are shown in Figure 6 for 4-day old rat pups in control and loperamide-treated groups. Qualitatively, methylene blue is seen to transit much less in the animals treated with loperamide as compared to vehicle-treated controls. Quantitative analysis of methylene blue transit velocity shows 8.73 +/- 0.86 cm/hr and 3.18 +/- 1.22 cm/hr in the control and loperamide-treated groups, respectively. This represents a 63.6% reduction in small intestinal transit velocity from loperamide administration as compared to control. These data confirm decreased intestinal motility in loperamide-treated animals as detected with US and PAI in Figure 5.

**Figure 6.**
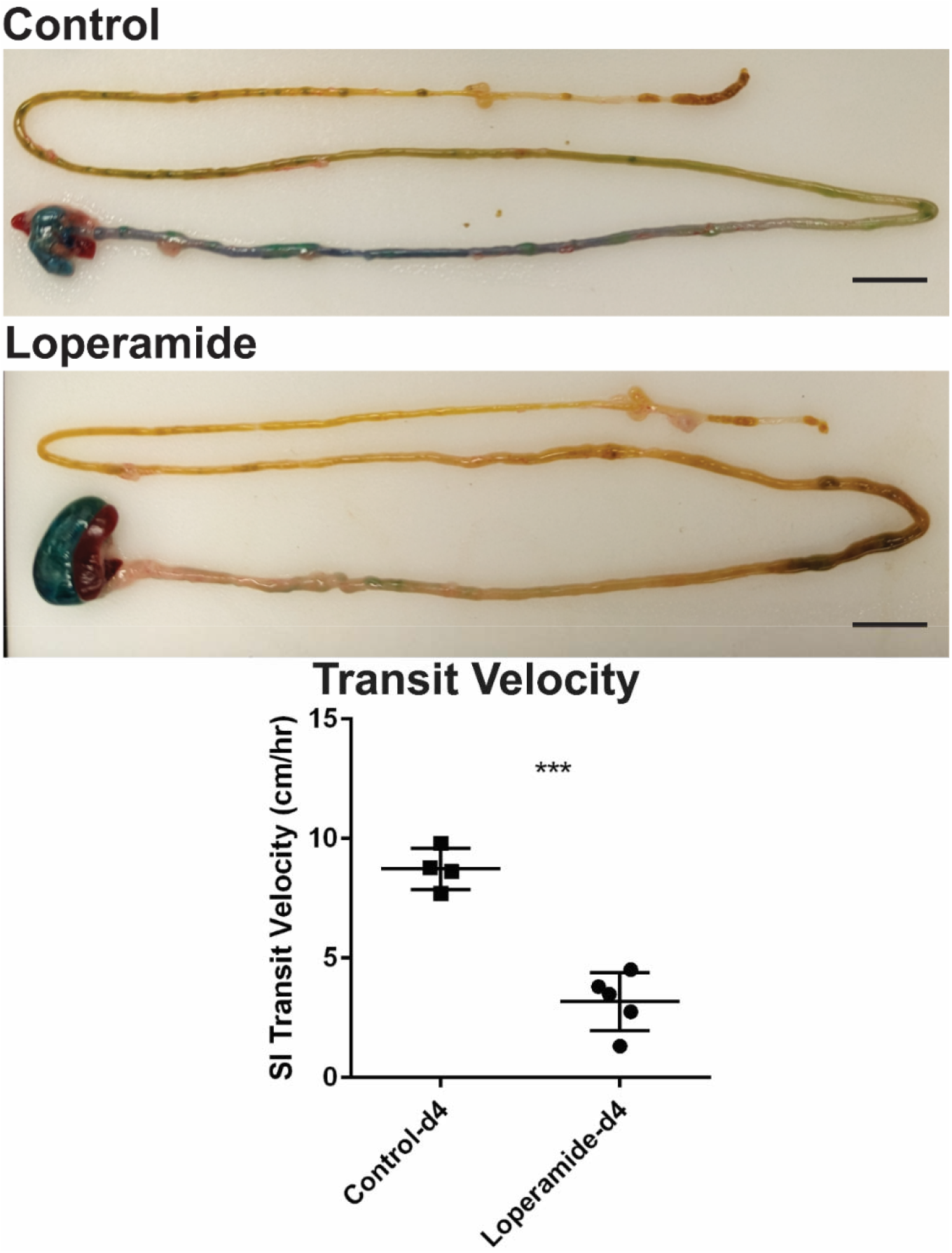
Small intestinal transit assay by methylene blue oral gavage in 4-day old control and loperamide-treated animals. The transit assay confirms a significant reduction in methylene blue transit with the graph demonstrating quantification of small intestinal transit velocity. (*** denotes p < 0.001. Scale bar = 1 cm).

## Discussion

Tissue-level blood oxygenation is known to dynamically change in response to alterations in FiO2 of inspired gas. Perturbation of FiO2 is an established method for validating PAI of oxygen saturation [21, 22]. In our study, we sought to examine changes in intestinal tissue oxygenation in response to stepwise changes in FiO2 through a breathing gas challenge comparing normoxia, hypoxia, and hyperoxia. Our results confirmed expectations in response to the breathing gas challenge with statistically significant differences in mean intestinal sO2 at each FiO2 level, 5%, 21%, and 100%. These results demonstrate that intestinal oxygenation imaging via PAI provides adequate sensitivity for differentiating intestinal tissue blood oxygenation in hypoxic, normoxic, and hyperoxic settings. These data validate the use of PAI to assess tissue oxygenation levels and dynamic changes within the intestine.

Loperamide is a widely used anti-diarrheal agent and inhibitor of intestinal peristalsis, acting as a mu-opioid receptor agonist with direct effects on intestinal smooth muscles to reduce peristalsis and act to increase overall intestinal transit time [23, 24]. We sought to develop a quantitative metric for non-invasively assessing motility from US and PAI imaging and examine changes in intestinal motility in response to loperamide administration. We adapted a previously established method based on assessing intestinal motility in humans from dynamic magnetic resonance imaging [20], and extended a similar analysis technique to assess intestinal motility from US and contrast enhanced PAI imaging in the neonatal rat intestine. Our results showed significant reduction in intestinal motility in response to loperamide administration in 4-day old rat pups, as expected. Reduction of intestinal motility in loperamide treated animals as compared to controls was confirmed by a physiological small intestinal transit assay with ex vivo measurement of methylene blue transit. These results demonstrate that intestinal motility assessment via contrast enhanced PAI and US imaging provides adequate sensitivity for detecting changes in intestinal motility, validating the use of PAI to assess intestinal motility.

In this study, we assessed intestinal tissue oxygenation and motility in healthy neonatal rats with acute perturbations under a breathing gas challenge and a drug administration study that transiently modified intestinal tissue oxygen saturation and intestinal motility, respectively, as a proof-of-concept. However, these transient perturbations do not entirely mimic the complicated disease processes typically found within the premature infant intestine. It will be important in future studies to further develop the use of PAI to measure intestinal tissue oxygenation and intestinal motility in intestinal disease onset and progression. One example is NEC disease, which is associated with compromised tissue oxygenation and intestinal dysmotility. Thus, NEC represents a promising next step for applying PAI. It will be important to evaluate PAI and analysis methods in an experimental animal model of NEC disease [25]. This would allow further in-depth paired analysis of PAI imaging with endpoint tissue histopathology. Additionally, our present study examines the utility of PAI for intestinal health and disease characterization in a preclinical animal study with rat pups at a developmental stage comparable to the intestine in premature human infants. This step is important to establish and validate imaging and analysis methods in the developing intestine under well-controlled experimental settings. However, this experimental design does not investigate potential regional variations of the intestine as found in disease states. As this technology is translated to the clinic for study in human infants, it will be important to incorporate assessment of regional heterogeneity in intestinal tissue oxygenation and motility. While the small physical size of the intestine of the neonate rat pup can present limitations on image resolution that may restrict full assessment of regional differences, future studies of PAI using the NEC rat pup model can develop and establish volumetric analysis methods of intestinal tissue oxygenation and motility within a disease model with regional variations. This volumetric analysis can be further optimized upon clinical translation into human infants to allow for more in-depth analysis of regional variation differences. Lastly, it is important to note that ultimate clinical application of current PAI technology for intestinal characterization will be likely limited to the infant/pediatric population due to physical imaging penetration depth limitations.

Tissue oxygenation and functional motility can be important markers for assessing the overall health and function of the intestine, but they are currently unable to be readily assessed in a premature infant. Due to their clinical fragility, the diagnostic imaging must be portable, non-invasive, and ideally with minimal radiation exposure. Thus, assessments are currently limited to portable XR which provides limited insight to tissue integrity. While US meets the necessary criteria for use in this population, it has been demonstrated to have variable utility due to lack of specificity and level of user/interpreter experience [14]. Our results show that the use of intestinal PAI can quantify changes in intestinal tissue oxygenation and intestinal motility, providing a potentially important adjuvant to existing diagnostic methods. Our results match the expected results from known experimental perturbations and demonstrate the sensitivity and resolution of PAI as a promising non-invasive imaging method for characterizing important biomarkers of physiology in the premature infant intestine. Additionally, these studies establish feasibility for the use of PAI with contrast agents that may be further developed to assess other physiological elements such as vascularization and intestinal uptake/permeability. Overall, these findings demonstrate the potential for a new non-invasive diagnostic method, with significant clinical translational potential, in assessing intestinal health and disease in premature infants.

In conclusion, assessing intestinal tissue oxygenation and motility are vital components to understanding intestinal physiology and pathology in premature infants. Imaging studies in this population have been limited to diagnostic XR and US, neither of which provide quantifiable measurements of tissue oxygenation nor motility. Diseases of the premature infant intestine are a leading cause of morbidity and mortality with limited diagnostic capabilities. Our proof-of-concept study is an important first step in investigating the potential for photoacoustic imaging to provide valuable non-invasive insight into intestinal health and disease to improve the care of premature infants.

## Abbreviations

PAI: (photoacoustic imaging),
FiO2: (fraction of inspired oxygen),
ICG: (indocyanine green),
NEC: (necrotizing enterocolitis),
ROI: (region-of-interest),
sO2: (oxygen saturation),
XR: (X-ray),
US: (ultrasound),
a.u.: (arbitrary units)

## Acknowledgements

These studies were supported by the Wake Forest Institute for Regenerative Medicine Pilot Award, NIH-NIDDK K01DK125633, American Gastroenterological Association Research Scholar Award in Health Disparities, and Wake Forest University School of Medicine Faculty Start-Up Funds. We acknowledge the Preclinical Ultrasound and Photoacoustic Imaging Core of Wake Forest University School of Medicine supported in part by the Wake Forest Clinical and Translational Science Institute (NIH NCATS UL1TR001420), NIH ORIP/OD-High End Instrumentation Grant S10 OD012330 and the Hypertension and Vascular Research Center.

## Statement of competing interests

The authors have no competing interests to declare.

## Supplementary materials

**Supplemental Figure 1.**
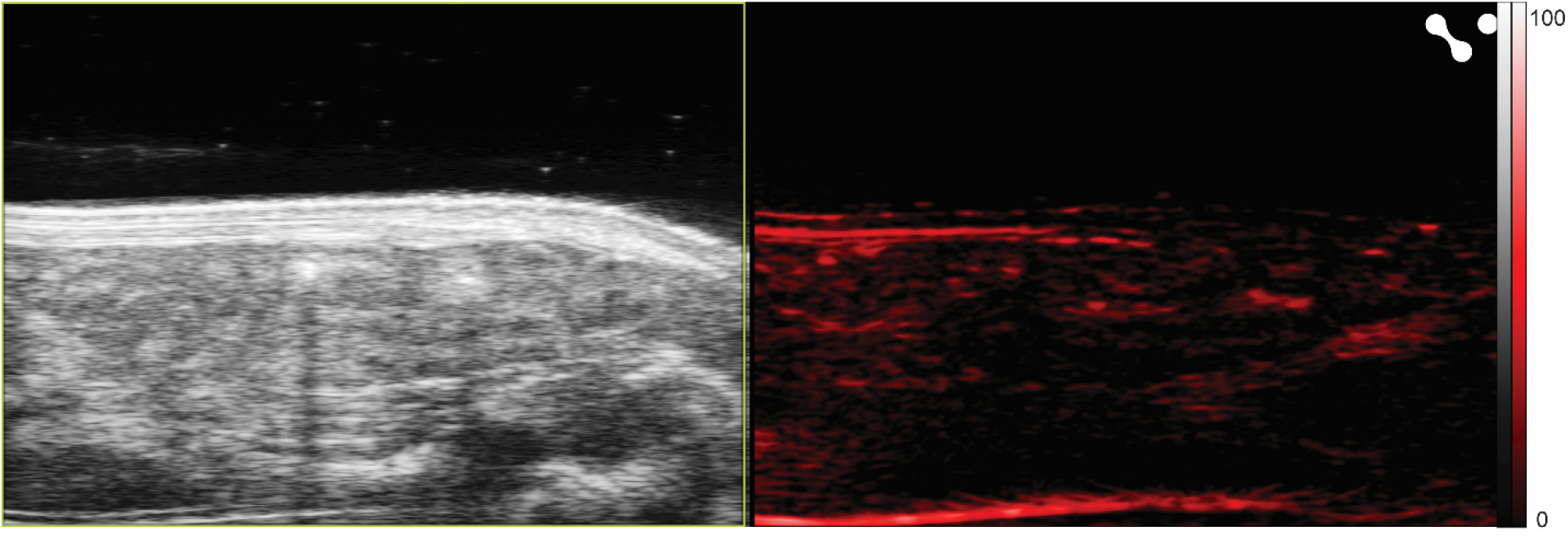
ICG-weighted PAI at 810 nm excitation wavelength in a 4-day old control rat pup without ICG contrast agent administration. Minimal intraluminal PAI signal is observed in an animal without oral gavage administration of ICG contrast agent when comparing to ICG-weighted PAI images in an animal with oral gavage administration of ICG contrast agent (see Figure 5). Residual ICG-weighted PAI signal at 810 nm excitation wavelength is due to background signal from hemoglobin.

**Supplemental Movie 1**. Cine time series movie (6 Hz frame rate) of intestinal peristalsis imaged by US and ICG-weighted contrast enhanced PAI in a 4-day old control rat pup.

**Supplemental Movie 2**. Cine time series movie (6 Hz frame rate) of intestinal peristalsis imaged by US and ICG-weighted contrast enhanced PAI in a 4-day old loperamide treated rat pup.

## Notes

### Competing Interest Statement

The authors have declared no competing interest.

